# Oxytocin, but not vasopressin, decreases willingness to harm others by promoting moral emotions of guilt and shame

**DOI:** 10.1101/2023.09.25.559242

**Authors:** Xiaoxiao Zheng, Jiayuan Wang, Xi Yang, Lei Xu, Benjamin Becker, Barbara J. Sahakian, Trevor W. Robbins, Keith M. Kendrick

## Abstract

Prosocial and moral behaviors have overlapping neural systems but whether they involve similar neurochemical systems is unclear. In the current pre-registered randomized placebo controlled trial on 180 adult male and female subjects we investigated effects of intranasal administration of two prosocial neuropeptides, oxytocin or vasopressin, on moral emotion ratings for situations involving accidental or intentional harm to others and in judgments of moral dilemmas where harm is inflicted for a greater good. Oxytocin, but not vasopressin, enhanced feelings of guilt and shame only for intentional harm and reduced endorsement of choices where direct intentional harm to others could achieve a greater good. Effects of oxytocin on guilt and shame were partially mediated by trait empathy. Overall, findings demonstrate for the first time that oxytocin, but not vasopressin, promotes unwillingness to deliberately harm others irrespective of the consequences. This may reflect stronger associations between oxytocin and empathy and vasopressin with aggression.

## Introduction

There is widespread interest in factors that influence the display of moral behavior given its key contribution to maintenance of stability in human social groups. Our capacity to exhibit prosocial behaviors is considered to be closely associated with that for making moral judgments and emotional responses to moral transgressions^1–4^, and there is considerable overlap between brain regions involved in social behavior and moral judgments and emotions^5,6,7^. There is also some support for this association between prosocial and moral behavior in disorders such as autism^8^ or psychopathy^9^ where reduced prosocial behaviors are paralleled to some degree by reduced moral behavior. However, it is still unclear the extent to which prosocial and moral behaviors are dissociable and one approach to addressing this is to determine whether neurochemical systems involved in social cognition and motivation are also important for moral judgments and associated emotional responses.

The majority of studies investigating the influence of neurochemical systems and moral behavior have focused on diffuse transmitter systems which influence a wide range of behaviors, primarily serotonin and dopamine. Serotonin in particular has been reported to influence moral judgments^10–13^ but also impacts social cognition and motivation as well as many other behaviors^14^. Similarly, dopamine may play a role in moral behavior decision making^12,15^ as well as in social motivation and reward^16^. However, to gain a greater potential insight into shared and distinct aspects of mechanisms contributing to prosocial and moral behavior it may be advantageous to consider neurochemical systems which have more selective effects on social and motivational behavior and determine whether they have similar or different effects on moral behavior.

In recent years, numerous studies have demonstrated key roles for the hypothalamic neuropeptides oxytocin (OXT) and vasopressin (AVP) and their receptors in social cognition and motivation in both animal models and humans^17,18^ and several clinical trials have reported improved social function in young children with autism after chronic intranasal treatment with both neuropeptides^19–22^. On the other hand, animal models in particular have demonstrated that the two peptides can produce opposite effects on stress and anxiety^23,24,25^ and aggression^26,27^.

While both OXT and AVP have extensive functional interactions with diffuse transmitters such as serotonin and dopamine which can influence moral behavior^28,29,30^, relatively little is known about their respective influences on moral behavior. Studies investigating effects of intranasal OXT in the social cognition domain most relevant to moral behavior have reported facilitation of emotion processing^31,32^, parochial and/or universal altruism^33,34^, emotional empathy^35,36,37^ and empathy for the pain of others^38^, paternal responses to children^39^, co-operative behavior^40–43^ and generosity^44^, ^45^as well as punishment for betrayal of trust^46^, although with some sex differences^43,46^. Intranasal OXT also produced sex-dependent effects in attraction to prospective partners who had previously been unfaithful in a relationship^47^. On the other hand, OXT can reduce jealousy and arousal in both sexes in response to either imagined or real partner infidelity^48^ and the impact of negative social feedback on learning^49,50^, suggesting that it may increase tolerance to moral transgressions and fear of social punishment. Several studies have investigated effects of intranasal OXT in the specific context of moral judgment tasks although with variable results. In an initial study OXT was reported to increase endorsement of moral dilemmas involving self-benefit in men but to decrease this in women^51^. However, another study in males found that OXT rendered individuals more forgiving of moral transgressions by others^52^. A further study in men found that OXT facilitated acceptance of all types of moral dilemmas faster than rejection and decreased orbitofrontal cortex responses for moral compared with non-moral dilemmas^53^. Some studies have reported that moral judgments are influenced by OXT receptor genotype^54,55,56^, although one focusing on utilitarian judgments did not^57^. Less is known about the effects of OXT on moral emotion responses to self- or other-actions, although one study has shown that it enhances self-embarrassment^37^.

Relatively few studies have been conducted on the vasopressin system. While, like OXT, it facilitates face emotion processing^58–61^, attention to social cues^62^ and the impact of negative social feedback^49^, it does not appear to influence altruism^34^, although two studies have reported some positive effects on co-operative behavior^43,63^. One study reported that AVP, in contrast to OXT, did not influence frontal and basal ganglia regions involved in empathy and reward processing when fathers looked at their children^39^, although another found that AVP relative to OXT evoked a small increase in empathic concern, but only in individuals who had received higher amounts of paternal warmth^64^. However, animal models have reported that while AVP facilitates male aggression^27^, OXT can reduce it^26^, and in humans intranasal AVP may promote aggressive reactions in males^65,66^ or pre-emptive defensive aggression in both sexes^67^. To date no studies have specifically investigated the effects of AVP on moral behavior.

From the above summary it is unclear the extent to which OXT and AVP influence moral judgments and emotional responses to moral transgressions and, if so, in which direction. The current study therefore investigated the effects of intranasal administration of both peptides on two different moral behavior tasks in a large cohort of adult male and female subjects (n = 180). In line with many studies on moral behavior^68,69^ both tasks focused on responses to scenarios involving deliberate harm to others. The first task uses scenarios with accidental or deliberate harm to others and the subject is required to report their emotional responses to each scenario imagining themselves to be either the agent or a victim (from the EMOTICOM battery^70^). The second task involves making decisions on whether to endorse actions in scenarios depicting personal or impersonal moral or non-moral dilemmas (from Greene et al 2001^71^). Here we specifically focused on responses to personal moral dilemmas involving willingness to directly physically harm one individual in order to save others. Given a background of more extensive evidence from previous studies for OXT influencing behaviors related to moral judgment, most notably empathy, we hypothesized that OXT would produce stronger effects than AVP on increasing feelings of shame and guilt for causing harm to others and on decreasing willingness to harm someone even when this could result in saving others. We also hypothesized that treatment effects on moral emotions and decisions might be influenced by how empathic individual subjects were in view of the well-known relationship between empathy and moral judgments^72^.

## Results

### Oxytocin, but not vasopressin, increases feelings of shame and guilt in the moral emotions task

We used 28 cartoons from the moral emotions task (EMOTICOM^70^) to examine the effects of OXT and AVP on emotion ratings when harming others as an agent or when exposed to harm as a victim respectively. 2-way repeated-measures ANOVAs were used with intention condition (deliberate harm/ accidental harm) as a within-subject factor and treatment (OXT/ AVP/ PLC) as a between-subject factor and ratings (agent - ashamed, guilty and feeling “bad”; victim - annoyed, feeling “bad”) served as dependent variables respectively.

For ratings of ashamed feelings, there was a significant intention condition ⅹ treatment interaction (F(2,159) = 5.762, *p* = 0.004, η_p_^2^ = 0.068). Post-hoc Bonferroni corrected tests showed no significant differences between treatments for ashamed ratings in the accidental harm condition (accidental harm: mean ± SD, OXT = 5.725 ± 0.903, PLC = 5.563 ±1.085, AVP = 5.739 ± 0.838; all *p*s > 0.992). However, in the context of causing deliberate harm to others, OXT significantly increased individuals’ ratings of ashamed feelings relative to both PLC and AVP (deliberate harm: mean ± SD, OXT = 5.409 ± 0.876, PLC = 4.844 ± 1.269, AVP = 4.912 ± 1.027; OXT vs PLC - *p* = 0.022, Cohen’s d = 0.517; OXT vs AVP - *p* = 0.049, Cohen’s d = 0.519); AVP and PLC did not differ from each other (*p* = 1.000) (Figure 1a).

Similarly, a significant intention condition ⅹ treatment interaction was observed for individual ratings of guilt (F(2,159) = 6.860, *p* = 0.001, η_p_^2^ = 0.079), due to the intention-specific effect of OXT on deliberate (deliberate harm: mean ± SD, OXT = 5.585 ± 0.829, PLC = 4.997 ± 1.221; AVP = 4.972 ± 0.996; OXY vs PLC - *p* = 0.012, Cohen’s d = 0.562; OXT vs AVP - *p* = 0.007, Cohen’s d = 0.666; AVP vs PLC - *p* = 1.000) rather than accidental harm (accidental harm: mean ± SD, OXT = 6.077 ± 0.741, PLC = 5.860 ±1.004, AVP = 6.035 ± 0.717; all *p*s > 0.545) (Figure 1b). Thus overall, OXT but not AVP, increased guilt and shame when participants imagined themselves as agents causing deliberate, but not accidental, harm to others.

**Figure 1.**
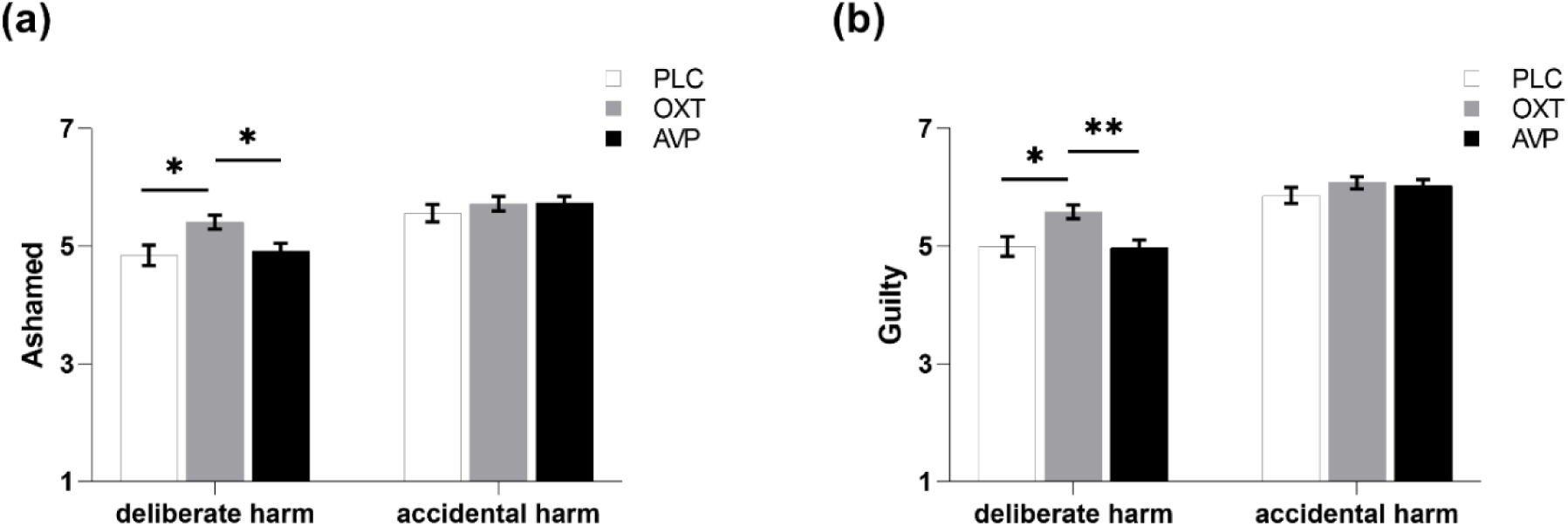
Intention conditionⅹdrug effect on moral emotions. OXT increased the ratings of feeling **(a)** ashamed and **(b)** guilty when deliberately harming others as an agent, relative to both PLC and AVP. * and ** denote significant Bonferroni-corrected post-hoc difference at *p* < 0.05 and *p* < 0.01 respectively. Bars represent means and SEM.

For ratings of feeling “bad”, there was no significant intention condition ⅹ treatment interaction (F(2,159) = 1.455, *p* = 0.236) or main effect of treatment (F(2,159) = 0.274, *p* = 0.761). However, a main effect of intention condition (F(1,159) = 206.196, *p* < 0.001, η_p_^2^ = 0.565) indicated that causing accidental harm as an agent was rated as feeling worse in comparison with causing deliberate harm (mean ± SD, deliberate harm = 2.699 ± 0.912, accidental harm = 1.878 ± 0.545, *p* < 0.001, Cohen’s d = 1.093) regardless of treatment. Thus overall, OXT but not AVP, specifically increased feelings of guilt and shame when participants imagined themselves as agents causing deliberate but not accidental harm to others.

For ratings by subjects as victims of harm there were no treatment-related effects for annoyance (intention condition ⅹ treatment interaction: F(2,159) = 0.407, *p* = 0.666; main effect of treatment: F(2,159) = 1.093, *p* = 0.338) or feeling “bad” (intention condition ⅹ treatment interaction: F(2,159) = 0.705, *p* = 0.495; main effect of treatment: F(2,159) = 0.084, *p* = 0.919). However, there were significant main effects of intention condition for both annoyance (F(1,159) = 209.62, *p* < 0.001, η_p_^2^ = 0.569) and feeling “bad” (F(1,159) = 215.40, *p* < 0.001, η_p_^2^ = 0.575) demonstrating that exposure to deliberate harm as a victim was rated more annoying (mean ± SD, deliberate harm = 5.711 ± 0.795, accidental harm = 4.828 ± 0.839; *p* < 0.001, Cohen’s d = 1.08) and made subjects feel worse (mean ± SD, deliberate harm = 1.765 ± 0.480, accidental harm = 2.377 ± 0.533; *p* < 0.001, Cohen’s d = 1.207) in comparison to accidental harm, irrespective of treatment.

### The effects of oxytocin on moral emotions are moderated by trait empathy

Based on the specific effects of OXT in the Moral Emotions Task, moderation analyses were performed to investigate moderation effects of three dimensions of empathy (perspective taking, fantasy and personal distress), on the behavioral indices which showed significant differences between OXT and PLC group. To this end treatment (OXT/ PLC) was used as the independent variable, ratings of guilt and ashamed feelings (when harming others deliberately as an agent) served as dependent variables respectively. Separate models were tested for each moderator (each of the three subscale scores of IRI here) and Bonferroni corrections (x6) were applied.

For personal distress, a dimension of empathy which refers to affective reactions in response to the experience of others, a significant moderation effect was found on both ashamed (R^2^ = 0.145, F(3,101) = 5.711, *p* = 0.001, *p*corrected = 0.007) and guilt (R^2^ = 0.172, F(3,101) = 7.007, *p* < 0.001, *p*corrected = 0.001) ratings, reflecting that scores on the personal distress dimension significantly moderated the effects of OXT on increasing ashamed (B = −0.136, S.E. = 0.046, *T*101 = −2.956, *p* = 0.004, *p*corrected = 0.023) and guilt ratings (B = −0.140, S.E. = 0.044, *T*101 = −3.222, *p* = 0.002, *p*corrected = 0.010; see Table S1) in the context of deliberately harming others. Further disentangling the effect using the Johnson-Neyman method^73^ revealed that only when the individual personal distress score was below 60% of the whole sample distribution (subscale score < 10.534; standardized z-score < 0.255) were treatment group differences on ashamed ratings as agents deliberately harming others considered significant (OXT > PLC, see Figure 2a), and only when the score was below 60% of the whole sample distribution (subscale score < 10.791; standardized z-score < 0.311), were treatment group differences on guilt ratings considered significant (OXT > PLC, see Figure 2b). No significant moderating effects were observed for the other subscales of IRI.

**Figure 2.**
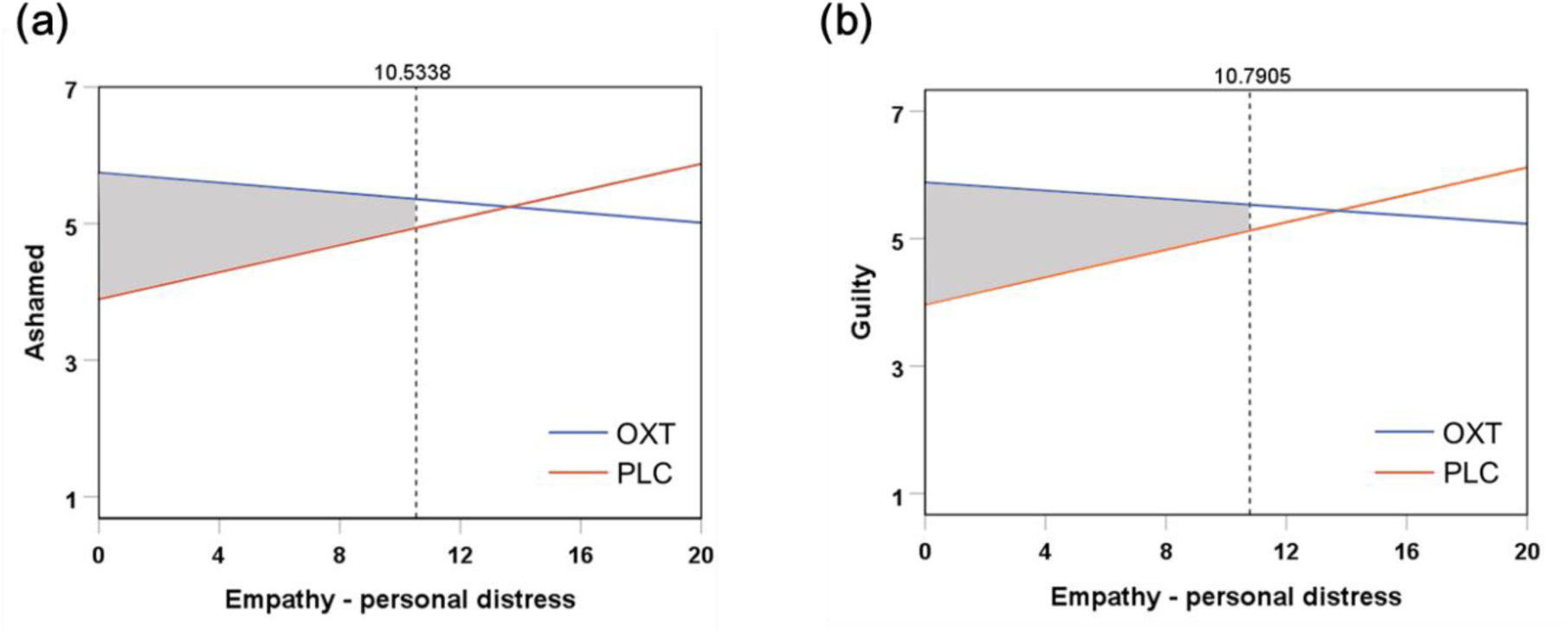
Individual specific empathy dimension - personal distress level modulated the OXT effect on ashamed and guilty feelings. OXT’s increased effect on ratings of **(a)** shame and **(b)** guilt in the context of harming others deliberately was evident when the individual personal distress was below 10.534 (60%; OXT > PLC) and 10.791 (60%; OXT > PLC) respectively. Shaded area identifies regions of significance in which ratings of feeling differ significantly between the OXT and PLC groups at *p* < 0.05. The outer border represents the lowest and highest score of empathy-personal distress subscale in the sample.

### Oxytocin, but not vasopressin, reduces willingness to endorse dilemmas involving deliberate harm in the moral judgment task

24 scenarios from Greene’s moral dilemma task^71^ were used to assess effects of OXT and AVP on moral judgments. A 2- way repeated-measures ANOVA was performed with endorsement rate (percentage) towards scenarios as the dependent variable, Greene’s taxonomy (non-moral/ moral impersonal/ moral personal) as a within-subject factor and treatment (OXT/ AVP/ PLC) as a between-subject factor.

Results yielded a significant taxonomy ⅹ treatment interaction (F(4,316) = 2.536, *p* = 0.043, η_p_^2^ = 0.031). Post-hoc Bonferroni corrected tests revealed that OXT selectively reduced endorsements compared with PLC towards moral personal scenarios (mean ± SD % endorsed, OXT = 49.28 ± 21.78%, AVP = 56.58 ± 18.31%, PLC = 59.86 ± 19.70%; OXT vs PLC - *p* = 0.023, Cohen’s d = 0.510; OXT vs AVP - *p* = 0.174; AVP vs PLC - *p* = 1.000) (Figure 3), while no effect of OXT or AVP was found for either moral impersonal (mean ± SD % endorsed, OXT = 63.94 ± 16.36%, AVP = 67.11 ± 15.96%; PLC = 68.51 ± 15.94%, all *ps* > 0.449) or non-moral (mean ± SD % endorsed, OXT = 83.65 ± 9.76%, AVP = 83.99 ± 9.67; PLC = 81.01 ± 11.74%, all *ps* > 0.412) scenarios, again implying an effect of OXT only on moral judgments in situations involving deliberate harm to others.

**Figure 3.**
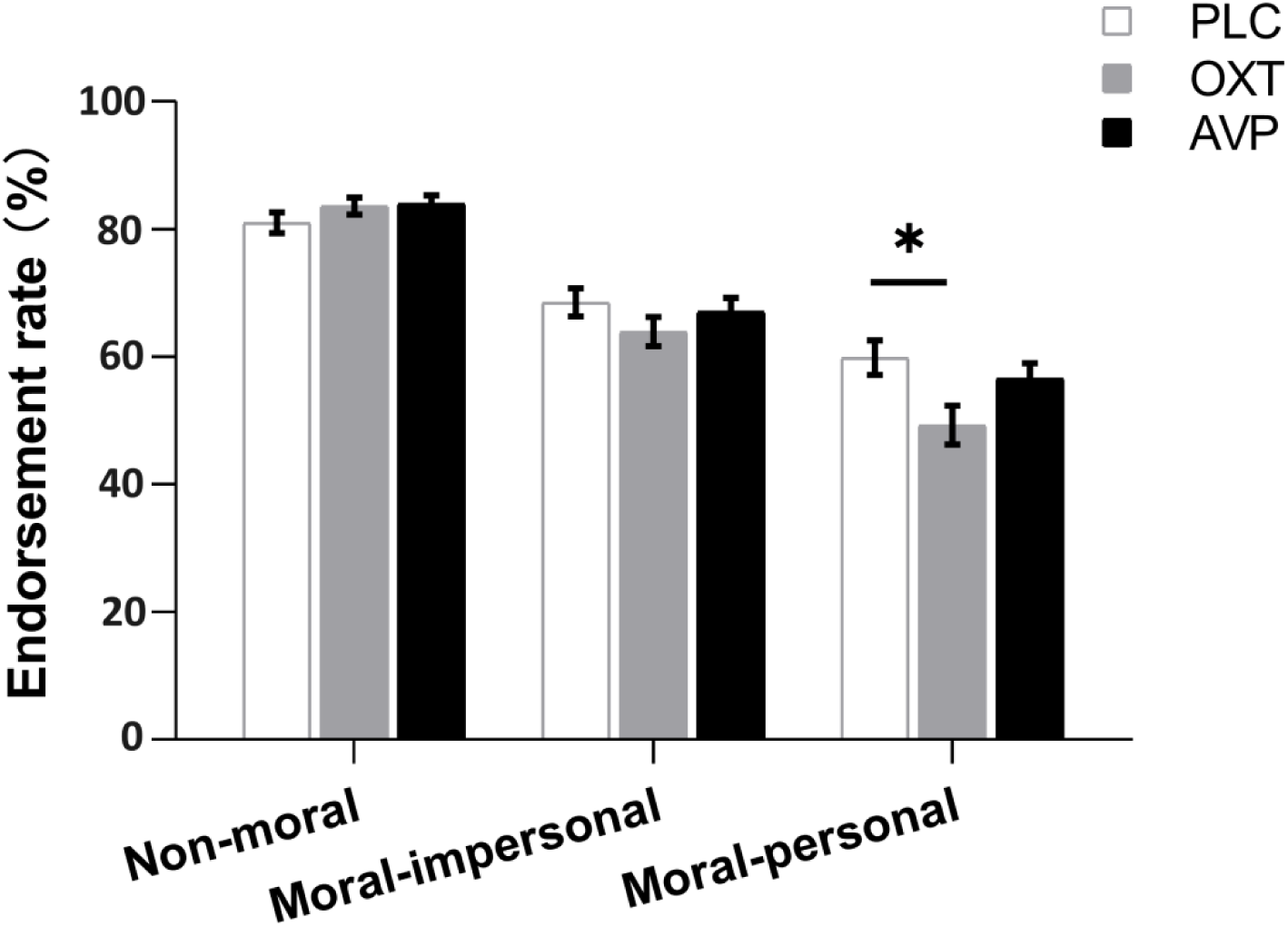
Greene’s taxonomy ⅹtreatment effect on moral judgment. OXT selectively reduced the endorsement rate towards moral personal scenarios in comparison to PLC. * denotes significant Bonferroni-corrected post-hoc difference at *p* < 0.05. Bars represent means and SEM.

For post-test arousal ratings for the different moral dilemmas an ANOVA revealed no significant taxonomy ⅹ treatment interaction (F(4,316) = 1.303, *p* = 0.275) or main effect of treatment (F(2,158) = 1.222, *p* = 0.297). However, there was a significant main effect of taxonomy (F(2,316) = 99.865, *p* < 0.001, η_p_^2^ = 0.387) indicating that general arousal responses towards the three scenario categories were different from one another, with arousal ratings for moral personal scenarios being the highest and those towards the non-moral scenarios being the lowest (mean ± SD, moral personal = 5.273 ± 1.323, moral impersonal = 4.425 ± 1.100, non-moral = 3.690 ± 1.216, all *p*s < 0.001). Thus, OXT specifically influenced endorsement of personal moral dilemmas involving directly inflicting personal harm but without altering general arousal associated with making decisions.

### Gender and age do not influence treatment-related effects

When both gender and age were included as covariates the results from remained robust (see supplementary Tables S2 and S3). A separate ANOVA analysis for the PLC group also revealed no significant gender differences in ratings and choices in the two tasks (see Table S4).

### Oxytocin and vasopressin reduce post-task negative mood but not positive mood or anxiety

To examine potential effects of treatment and the task on mood (both positive and negative PANAS subscales) and state anxiety (SAI), 2-way ANOVAs with time point (pre-test/ post-test) as within subject factor and drug (OXT/ AVP/ PLC) as between subject factor were further conducted. Results demonstrated a significant time point ⅹ treatment interaction (F(2,158) = 3.410, *p* = 0.035, η_p_^2^ = 0.041) for negative mood, due to reduced negative feelings after both OXT (mean ± SD, pre-test = 19.962 ± 6.666; post-test = 17.923 ± 6.642; *p* = 0.002, Cohen’s d = 0.306) and AVP (mean ± SD, pre-test = 18.614 ± 6.646; post-test = 16.842 ± 6.976; *p* = 0.006, Cohen’s d = 0.260) treatment but not in the PLC group (mean ± SD, pre-test = 17.788 ± 5.856; post-test = 17.981 ± 7.103; *p* = 0.772). No significant time point ⅹ treatment interactions were found for positive mood (F(2,158) = 2.510, *p* = 0.085) or anxiety scores (F(2,158) = 0.049, *p* = 0.952). One female subject in the PLC group was excluded in this analysis since post-test responses were not collected.

## Discussion

The current study investigated the roles of the two closely related hypothalamic neuropeptides AVP and OXT on moral emotions and judgments using two different paradigms. In support of our original hypotheses, we have demonstrated for the first time that only intranasal OXT administration significantly influenced moral emotions with it specifically increasing feelings of shame and guilt when causing deliberate harm to others but without generally influencing how bad or annoyed individuals felt. In further support of this we demonstrated that OXT specifically reduced endorsements of moral personal choices involving deliberate harm to anyone, no matter what the justification, but without influencing individuals’ general arousal levels. Importantly, OXT had no effects on feelings of shame and guilt as a result of accidentally harming others or on endorsing impersonal or non-moral moral dilemmas. Furthermore, OXT had no effects on emotional ratings when participants imagined themselves as victims of harm as opposed to agents inflicting it.

Overall, the findings from responses to both moral judgment tasks in the current study are firstly that intranasal OXT specifically increases feelings of the self-conscious moral emotions, guilt and shame, when an individual imagines causing deliberate harm to others, but without influencing more general feelings (i.e. feeling bad or annoyed). Secondly, OXT specifically decreases the likelihood that individuals will endorse inflicting any deliberate harm to others, no matter what the beneficial consequence of their actions might be, and without influencing general arousal. On the other hand, OXT had no influence on feelings of guilt and shame when harm towards others was accidental or on endorsements of decisions which might indirectly cause harm to specific individuals but save others. As such, OXT may be acting to strengthen deontological moral decision making where it is the morality of the action performed rather than its consequence which is important. It also corresponds to the principle of non-maleficence proposed by Beauchamp and Childress^74^ in the context of biomedical ethics. This facilitation of deliberate harm aversion following OXT treatment is very much in line with its well established prosocial and empathic effects ^17,18^ and also the strong association between moral and social behaviors^1–4^. Although sex differences have been reported for the influence of OXT on some aspects of social cognition and sharing^75,76^ and judgments of others’ positive and negative characteristics^77^ we found no evidence for this for its effects on moral emotions or judgments. While one previous study found opposite effects of OXT on endorsement of self-benefit moral decisions in males and females^51^, in a subsequent study no gender differences were found and OXT increased the speed with which moral dilemmas were endorsed^53^. Thus, it is possible that there may be some sex differences on the effects of OXT on certain kinds of moral dilemmas.

A moderation analysis of the influence of trait empathy (IRI sub-scales) on the effects of OXT on moral emotions ratings revealed that they were moderated by scores on the personal distress sub-scale. Thus, individuals scoring low on the personal distress scale were more likely to exhibit OXT-mediated increases in ratings of shame and guilt. Personal distress is a measure of the affective response to the experiences of others and so OXT appears to mainly facilitate increased harm avoidance in individuals with lower emotional responses to the suffering of others, although only when such suffering is caused by them. Overall, this is in line with our previous findings that intranasal OXT facilitates emotional empathy in both men and women which may be mediated via its actions on the amygdala^35,36,37,78^ and other findings that it can enhance feelings of empathy towards members of an out-group experiencing pain^38^.

How might OXT be promoting increased aversion to deliberately harming others? Findings from neuroimaging^7,79,80^, brain stimulation^81^ and brain lesion^82^ studies have consistently implicated the medial frontal cortex and is limbic connections with moral emotions and decision making. Intranasal OXT strengthens functional connectivity between the medial prefrontal cortex and limbic regions^83,84,85^ and OXT receptor mRNA is decreased in the medial prefrontal cortex in a number of psychiatric disorders where moral behavior can be affected^86^. Oxytocin has extensive functional interactions with other transmitter systems^28,29^, and one possibility in the current moral behavioral context is through its interactions with serotonin signaling in the brain. Increasing serotonin concentrations using serotonin selective re-uptake inhibitor (SSRI) drugs such as citalopram increases OXT release^87^, and has been shown to increase harm aversion in a several experimental contexts, including moral decision making and unfair behavior in economic games^10–13^. Intranasal OXT also alters serotonin receptor activity (5HT1-A) in a number of frontal cortex and limbic brain regions associated with moral decision making^88^. Furthermore, the basal ganglia effects of OXT on social reward behavior in animals^89^ and its amygdala effects during threat processing in humans^90^ are dependent to some extent on serotonin signaling. While established OXT interactions with dopamine signaling^91^ might also play a role, a study has shown that in contrast to serotonin increasing dopamine function does not influence harm aversion per se but has more of an influence on altruism^12^.

Given the closeness of AVP and OXT to one another in terms of structure and receptor actions, as well as evidence that they can produce similar neural and behavioral effects in social contexts such as face emotion processing^58–61^, attention to social cues^62^ and co-operative behavior^43,63^, it is perhaps surprising that in the current study AVP had no significant effect on moral emotions or decision making in the two tasks we used. However, there is less support for an involvement of AVP in behaviors related to moral behavior such as altruism and empathy and more on risk taking and both offensive and defensive aggression^27,65,66,67^, particularly in males^92^. In the context of empathy, it should be noted that while one study found no evidence that AVP altered neural responses in brain regions involved in empathic responses when fathers viewed their children^39^, another study has reported that AVP, but not OXT, increased empathic concern, although only in the context of individuals receiving high levels of paternal warmth^64^. However, in contrast to OXT, to date no studies have reported general effects of AVP on emotional empathy. Furthermore, although AVP also interacts with serotonin systems, animal studies suggest this may be more antagonistic, with serotonin blocking effects of AVP on offensive aggression^93^ and SSRIs reducing rather than increasing brain AVP^94^. Thus, while AVP like OXT may act to promote a number of pro-social behaviors, only OXT appears to facilitate moral behaviors in terms of harm aversion, possibly reflecting AVP’s more prominent role in offensive aggression. Interestingly though, we found that both OXT and AVP reduced post-task negative mood scores compared to PLC.

Several limitations should be acknowledged in the current study. Firstly, only a single dose of AVP and OXT were used and possibly AVP might have influenced moral emotions and decision making at higher doses. However, the doses used of the two peptides were equimolar and a 20IU AVP dose has been reported to have both behavioral and neural effects by a number of different studies^43,62,64,65,66^. Indeed, we also found that both OXT and AVP reduced post-task negative mood scores in the current study. Secondly, moral emotions and decisions were only made in imagined rather than real-life contexts.

## Conclusions

Our findings demonstrate for the first time that the neuropeptide OXT specifically enhances feelings of shame and guilt when imagining causing deliberate harming to others and correspondingly reduces the likelihood that individuals will choose to do so even when the outcome might save others. These specific effects of OXT on harm aversion were not observed following treatment with the closely related neuropeptide AVP. Thus, despite having many similar prosocial effects the two peptides differ markedly in their modulation of moral behavior demonstrating dissociable effects on the neurochemical control of prosocial and moral behaviors. This may possibly reflect a greater role for OXT in promoting altruism and empathy and for AVP in promoting aggression.

## Materials and Methods

### Participants

180 healthy adults (90 males, 18-26 years, mean ± SD age = 20.53 ± 1.880 years) were recruited from the University of Electronic Science and Technology of China (UESTC). All participants were free from medical or psychiatric disorders and were instructed to abstain from alcohol, caffeine, nicotine, or medications during the 24 h before the experiment (self-report). Female participants were not pregnant or in their menstrual period or taking oral contraceptives (self-report). The protocol was approved by the local Ethics Committee of the UESTC (number 101420210426008) and adhered to the latest revision of the Declaration of Helsinki. Participants provided written informed consent before the start of the experiment and were financially compensated after participation (110 RMB). The study was pre-registered as a clinical trial (https://clinicaltrials.gov/show/NCT04890470).

### Experimental procedure

To control for the potential between-group differences in verbal IQ, mood, clinical symptoms, personality traits, ethical attitudes, a batch of validated Chinese questionnaires were implemented before the drug application (see supplementary materials). The Positive and Negative Affective Scale - PANAS^95^ and State and Trait Anxiety Inventory - STAI^96^ were administered before treatment and after completion of the tasks to measure treatment/task effects on mood and state anxiety. Preliminary one-way ANOVAs were implemented to examine group differences in age, verbal IQ, moral values, psychological and personality traits among the OXT-, AVP- and PLC-treated subjects and showed no significant group differences (see supplementary Table S5). The Interpersonal Reactivity Index - IRI^97^ was administered prior to treatment to permit associations between task performance and trait empathy to be made. Participants were randomized to receive either 24 international units (IU) OXT, 20 IU AVP or PLC 45 min prior to the start of two experimental tasks involving moral decision making. During this 45-minute period participants could read either technical magazines or play solitaire card games on a computer to avoid evoking any form of competitive aggression stimulation. Next, subjects completed the Moral Emotions task and the Moral Judgment task (presentation order of both tasks was balanced).

### Treatment administration

In the current double-blind, between-subject, placebo-controlled design experiment, all participants were randomly allocated into OXT (n = 60, 31 males, mean ± SD age = 20.550 ± 1.845 years), AVP (n = 60, 30 males, mean ± SD age = 20.133 ± 1.799 years) or PLC group (n = 60, 29 males, mean ± SD age = 20.900 ± 1.946 years). A 24 IU OXT dose (OXT spay supplied sterile by Sichuan Defeng Pharmaceutical Co., Ltd, China; six 0.1ml puffs, 3 to each nostril and interspaced by 30 s) and 20 IU AVP dose (AVP supplied by Bio-Techne China Co., Ltd and was administered dissolved in identical ingredients of sodium chloride and glycerol as OXT nasal spray; 0.22μm Millipore filtered - six 0.1ml puffs, 3 to each nostril and interspaced by 30 s) were chosen since they are equivalent (i.e. equimolar) given the different molecular weights of the peptides. Indeed, the same 20 IU AVP dose has also been adopted in multiple previous studies^43,62,64^. The sterile PLC spray was identical in composition other than the neuropeptide (supplied sterile by Sichuan Defeng Pharmaceutical Co., Ltd, China). Intranasal treatment was self-administered by participants 45 min before the start of the experimental tasks following a standardized protocol^98^.

### Moral Emotions Task

The Moral Emotions task was based on the one in the neuropsychological test battery EMOTICOM^70^ but re-programmed via PsychoPy2^99^ with text in Chinese. Experimental stimuli were 14 different cartoon figures depicting moral scenarios with half involving a deliberate harm and half an accidental harm to others. Cartoon figures were edited to appear more Asian by giving them all black hair and brown eyes. Each cartoon figure was presented twice in the task and subjects were asked to imagine how they would feel in the situation as either the agent or the victim and to give corresponding ratings. Thus, during the Moral Emotions task a total of 28 cartoon scenarios were displayed randomly to each participant and feelings of ashamed, guilty, and feeling “bad” were rated when the subject imagined themselves as the agent, while ratings of annoyed and feelings “bad” were given when imagining themselves as a victim. A 7-point Linkert scale was adopted in this task (ashamed, guilty and annoyed: 1= not at all and 7 = extremely; feeling “bad”: 1 = feeling bad and 7 = feeling good). There was no time limitation for watching the cartoons or giving ratings during the task (Figure S1).

### Moral Judgment Task

Twenty-four hypothetical descriptions in Chinese depicting non-moral, moral indirect personal harm, and moral direct personal harm scenarios (8 scenarios per condition) were selected and adopted in the current task via a pilot study involving 40 (n = 20 female) independent subjects (details see Support Information: Materials and Methods) from the original 64 scenarios in Greene et al.^71^. Both the moral categories involved scenarios where the subject is given a choice of endorsing a utilitarian option which saves more compared with less people from harm. In the moral direct personal dilemma scenarios the subject as an agent is asked if they will endorse actions involving causing direct harm to individuals in order to save more whereas for the moral indirect impersonal dilemmas the subject agent is asked if they will endorse indirect actions which will result in saving more individuals from harm. A set of non-moral scenarios were also selected as control stimuli. The selected 24 scenarios were correctly classified as representing the specific categories (mean accuracy = 76.35%) and matched for endorsement rate and arousal ratings (see Support Information). Examples of scenarios are provided in Support Information Table S6.

The Moral Judgment task was programmed via E-prime 2.0 (http://www.pstnet.com/eprime.cfm, Psychology Software Tools, USA). All 24 verbal descriptions were randomly displayed for passive viewing and subsequently paired with a posed question relevant to the current scenario (“Is it appropriate to…?”). Subjects needed to make forced-choice decisions of either “yes” or “no” by pressing designated keys. For the two types of moral scenarios, endorsement refers to a utilitarianism choice and rejection refers to a deontological choice. There was no time limitation for reading the scenarios or responding to the posed question (Figure S2). At the end of the experiment the subjects were presented with all the moral dilemmas again and asked to rate how aroused they felt by them using a Likert scale (from 1 not at all to 7 extremely aroused).

### Statistical Analysis

A total of 18 subjects included in the initial randomization had to be excluded (8 in the OXT group, 3 in the AVP group and 7 in the PLC group), due to either technical problems with collecting the rating data in both experiments (n = 11), failing to understand instructions properly (n = 3) or to answer sufficient number of the questions (n = 2), having a cold (n =1), or revealing that they had taken medication in the last 24h (n = 1). These exclusions resulted in 162 subjects being included in the final analysis (52 subjects (26 males) in the OXT group, 57 subjects (29 males) in the AVP group and 53 subjects (26 males) in the PLC-control group). One additional subject had to be excluded only for the Moral Judgment task, due to a data collection technical failure (see Consolidated Standards for Reporting of Trials (CONSORT) flow diagram in Supplementary, Figure S3). Subjects who completed the study (n = 162) could not identify which treatment they received better than chance (33.3%) after the experiment (48 subjects guessed correctly, χ^2^ = 1.000, *p* = 0.317), confirming successful double-blinding.

All statistical analyses were performed using SPSS 26.0 software (SPSS Inc., Chicago, Illinois, USA). Based on the previous studies exhibiting gender differences for moral judgments, gender effects were firstly explored using repeated-measures ANOVAs only in the PLC group (see Supplementary and Table S2). Treatment effects were analyzed primarily using repeated-measures ANOVAs (individual factor details provided in the following Results section) and where significant interactions occurred post-hoc analyses were carried out with a Bonferroni correction. For both ANOVAs and post-hoc tests measures of effect size are also provided (Partial eta squared (η_p_^2^) or Cohen’s d).

For the behavioral indices that showed significant differences between treatment (OXT or AVP) and PLC-control groups, we further explored whether treatment effects would be moderated by trait empathy. Four dimensions of empathy were assessed by the IRI scale, including perspective taking (PT), fantasy (FS), empathic concern (EC), and personal distress (PD) Only three of them exhibited acceptable internal consistency in the current sample and served as moderators (Cronbach’s α scores: 0.78 for perspective taking, 0.69 for fantasy, and 0.778 for personal distress). Empathic concern only achieved an α of 0.58 and thus was excluded for further analyses. Separate models were tested for each moderator, as well as each behavioral outcome. We estimated the moderation effects using the PROCESS macro for SPSS (Model 1)^100^. Bonferroni corrections were applied for multiple comparisons. The Johnson-Neyman method^73^ was additionally used to calculate the entire range of moderator variable values (i.e., personal distress subscale scores) for which the focal predictor (i.e., treatment groups- OXT/PLC) is significantly (a = 0.05) associated with the dependent variable (i.e., ratings of ashamed feelings).

Gender and age were further subjected as covariates for the above-mentioned ANOVAs and moderation analyses to determine the potential effects of them on treatment-related effects on moral emotions and endorsement.

## Acknowledgements

The work was funded by National Natural Science Foundation of China (NSFC) [grant number 31530032], Key Scientific and Technological projects of Guangdong Province [grant number 2018B030335001] and UESTC High-end Expert Project Development grant [grant number Y0301902610100201]. Any opinions, findings, conclusions or recommendations expressed in this publication (or by members of this State Key Laboratory) do not reflect the views of the Government of the Hong Kong Special Administrative Region or the Innovation and Technology Commission

## Author contributions

XZ, TR & KMK designed the study; XZ, JW, XY and LX collected the data; XZ, LX and KMK performed the analysis. XZ & KMK wrote the first draft. XZ, BB, BJS, TR & KMK contributed to revised drafts. All authors read and approved the final submission.

