## Supplementary for "Oxytocin, but not vasopressin, decreases willingness to harm others by promoting moral emotions of guilt and shame"

### Supplementary Materials and Methods

**Scales and Questionnaires.** A batch of validated Chinese scales/questionnaires were implemented measuring verbal IQ, mood, clinical symptoms, moral attitudes and traits before subjects self-conducted the intranasal treatment to control for potential inter-group confounders. These included: the vocabulary test of Wechsler Adult Intelligence Scale Chinese Revised (WAIS-RC)^1^; State and Trait Anxiety Inventory - STAI^2^; Beck’s Depression Inventory - BDI^3^; Autism Spectrum Quotient - ASQ^4^;  NEO Five - Factor Inventory - NEO-FFI^5^; Childhood Trauma Questionnaire - CTQ^6^; Parenting Style Scale - EMBU^7^; Ethics Position Questionnaire - EPQ^8,9^; The Selfishness Questionnaire - SQ^10^; and Individualism-Collectivism Scale - ICS^11^. The Interpersonal Reactivity Index - IRI^12^ was also given to investigate associations between behavioral findings and empathy. The Positive and Negative Affective Scale - PANAS^13^ was completed before treatment and after the task to measure treatment and task-dependent changes in mood. The STAI (only state anxiety) questionnaire was also completed for a second time after the task to assess any treatment/task dependent effects on anxiety.

**Chinese Translations of the two tasks.** To create Chinese versions of the Moral Emotion and Moral Judgment tasks, the English was first translated into Mandarin Chinese by a bilingual expert and subsequently back-translated by another bilingual expert. The final translation was then checked by a native English speaker. Modifications were made on the Chinese drafts of the translation to eliminate all discrepancies between the original versions and the back-translated ones.

**Supplementary results**

**Pilot study of materials for the Moral Judgment Task.** N = 40 university subjects (20 males, mean ± SD age = 20.730 ± 1.519 years) participated in the pilot study to initially rate the Chinese version of the original 64 scenarios (25 for moral-personal; 19 for moral-impersonal; 20 for non-moral) from Greene et al^14^. General arousal (7 point Likert scale, 1 = not at all and 7 = extremely intense) and moral judgment (endorse or reject) towards each scenario, and accuracy of category discrimination were acquired and finally 24 scenarios were selected (8 scenarios for each condition) after excluding those which where endorsed by all subjects or deemed inappropriate for them and then balanced for the same number in each condition.

A two-way ANOVA with gender as a between-subject factor, Greene’s taxonomy (non-moral/ moral impersonal/ moral personal) as a within-subject factor, and endorsement rate as dependent variable was conducted. No significant gender ⅹ taxonomy interaction (F_(2,76)_ = 1.557, *p* = 0.219) or main effect of gender (F_(1,38)_ = 3.323, *p* = 0.076) was found. However, there was a significant main effect of taxonomy (F_(2,76)_ = 67.675, *p* < 0.001, η_p_^2^ = 0.640) indicating that the endorsement rates towards the three scenario categories were different from one another, with the endorsement rate for moral personal scenarios being the lowest and those towards the non-moral scenarios being the highest (mean ± SD % endorsed, moral personal = 50.31 ± 20.51%, moral impersonal = 58.75 ± 18.17%, non-moral = 86.87 ± 9.37%, all *p*s < 0.016).

For arousal ratings, the same ANOVA was implemented and no significant gender ⅹ taxonomy interaction (F_(2,76)_ = 3.629, *p* = 0.054) or main effect of gender (F_(1,38)_ = 0.126, *p* = 0.724) was found. There was also a significant main effect of taxonomy (F_(2,76)_ = 169.017, *p* < 0.001, η_p_^2^ = 0.816), indicating that moral personal scenarios were rated as the most arousing and non-moral ones rated as the least arousing (mean ± SD, moral personal = 5.738 ± 0.930; moral impersonal = 4.606 ± 0.986; non-moral = 2.669± 1.076, all *p*s < 0.001).

**Gender differences irrespective of treatment**

**Moral Emotions Task.** To explore gender differences for moral emotions irrespective of treatment, 2-way repeated-measures ANOVAs were carried out with intention condition (deliberate harm/ accidental harm) as a within-subject factor and gender (male/ female) as a between-subject factor and ratings (agent - ashamed, guilty and feeling “bad”; victim - annoyed, feeling “bad”) as dependent variables respectively, only for the PLC group. Results showed no intention condition ⅹ gender interaction (all *p*s > 0.099) or main effect of gender (all *p*s > 0.060) (see Table S2), suggesting no gender differences on the moral emotions task.

**Moral Judgment Task.** Similarly, to explore gender differences for moral judgment regardless of the treatment effect, a 2-way repeated-measures ANOVA with Greene’s taxonomy (non-moral/ moral impersonal/ moral personal) as a within-subject factor and gender (male/ female) as a between-subject factor and endorsement rate as a dependent variable only for the PLC group. Results showed no significant taxonomy ⅹ gender interaction (*p* = 0.224) or main effect of gender (*p* = 0.380) (see Table S2), indicating no gender differences on the moral judgment task.

**Table S1.** Results of moderation analyses in the Moral Emotions task.

| **Dependent variables** | **B** | **S.E** | **t** | ***p*** | ***p_corrected_*** | **R^2^** | **F** | ***p*** | ***p_corrected_*** |
| --- | --- | --- | --- | --- | --- | --- | --- | --- | --- |
| ***Ashamed feelings in the context of harming others deliberately as an agent*** | | | | | | | | |  |
| Treatment | 1.8495 | 0.4780 | 3.8694 | 0.0002 | 0.0012 | 0.1450 | 5.7105 | 0.0012 | 0.0072 |
| Personal distress | 0.0990 | 0.0349 | 2.8398 | 0.0055 | 0.0330 |  |  |  |  |
| Interaction | -0.1356 | 0.0459 | -2.9557 | 0.0039 | 0.0234 |  |  |  |  |
| ***Guilty feelings in the context of harming others deliberately as an agent*** | | | | | | | | |  |
| Treatment | 1.9159 | 0.4529 | 4.2307 | 0.0001 | 0.0006 | 0.1723 | 7.0070 | 0.0002 | 0.0012 |
| Personal distress | 0.1076 | 0.0330 | 3.2563 | 0.0015 | 0.0090 |  |  |  |  |
| Interaction | -0.1400 | 0.0435 | -3.2215 | 0.0017 | 0.0102 |  |  |  |  |

Notes: *p*_corrected_ – Bonferroni correction (x6).

**Table S2.** Results of the ANCOVA analysis after controlling for age and gender.

| **Dependent variable** | **ANOVA results** |
| --- | --- |
| ***Moral Emotions Task*** | |
| ***Agent – ashamed*** | |
| IntentionⅹTreatment | (F_(2,157)_ = 5.672, *p* = 0.004, η_p_^2^ = 0.067 |
| Intention | (F_(1,157)_ = 1.970, *p* = 0.162 |
| Treatment | (F_(2,157)_ = 2.015, *p* = 0.137 |
| ***Agent – guilty*** | |
| IntentionⅹTreatment | (F_(2,157)_ = 6.844, *p* = 0.001, η_p_^2^ = 0.080 |
| Intention | (F_(1,157)_ = 1.274, *p* = 0.261 |
| Treatment | (F_(2,157)_ = 3.429, *p* = 0.035, η_p_^2^ = 0.042 |
| ***Moral Judgment Task*** | |
| ***Endorsement rate*** | |
| TaxonomyⅹTreatment | (F_(4,312)_ = 2.787, *p* = 0.029, η_p_^2^ = 0.034 |
| Taxonomy | (F_(2,312)_ = 1.533, *p* = 0.218 |
| Treatment | (F_(2,156)_ = 2.618, *p* = 0.076 |

**Table S3.** Results of moderation analyses in the Moral Emotions task after controlling for age and gender.

| **Dependent variables** | **B** | **S.E** | **t** | ***p*** | ***p_corrected_*** | **R^2^** | **F** | ***p*** | ***p_corrected_*** |
| --- | --- | --- | --- | --- | --- | --- | --- | --- | --- |
| ***Ashamed feelings in the context of harming others deliberately as an agent*** | | | | | | | | |  |
| Treatment | 1.7680 | 0.4807 | 3.6782 | 0.0004 | 0.0024 | 0.1636 | 3.8720 | 0.0030 | 0.0180 |
| Personal distress | 0.0859 | 0.0367 | 2.3412 | 0.0212 | 0.1272 |  |  |  |  |
| Interaction | -0.1308 | 0.0460 | -2.8464 | 0.0054 | 0.0324 |  |  |  |  |
| ***Guilty feelings in the context of harming others deliberately as an agent*** | | | | | | | | |  |
| Treatment | 1.8155 | 0.4500 | 4.0347 | 0.0001 | 0.0006 | 0.2094 | 5.2424 | 0.0003 | 0.0018 |
| Personal distress | 0.0850 | 0.0344 | 2.4749 | 0.0150 | 0.0900 |  |  |  |  |
| Interaction | -0.1329 | 0.0430 | -3.0876 | 0.0026 | 0.0156 |  |  |  |  |

Notes: *p*_corrected_ – Bonferroni correction (x6).

**Table S4**. Results of ANOVA investigating gender differences for Moral Emotions and Moral Judgment Tasks for the PLC group.

| **Dependent variable** | **ANOVA results** |
| --- | --- |
| ***Moral Emotions Task*** | |
| ***Agent – ashamed*** | |
| Intention ⅹ Gender | (F_(1,51)_ = 0.120, *p* = 0.730 |
| Intention | (F_(1,51)_ = 33.314, *p* < 0.001, η_p_^2^ = 0.395 |
| Gender | (F_(1,51)_ = 2.526, *p* = 0.118 |
| ***Agent – guilty*** | |
| Intention ⅹ Gender | (F_(1,51)_ = 2.821, *p* = 0.099 |
| Intention | (F_(1,51)_ = 43.300, *p* < 0.001, η_p_^2^ = 0.459 |
| Gender | (F_(1,51)_ = 3.690, *p* = 0.060 |
| ***Agent – feeling “bad”*** | |
| Intention ⅹ Gender | (F_(1,51)_ = 1.680, *p* = 0.201 |
| Intention | (F_(1,51)_ = 65.924, *p* < 0.001, η_p_^2^ = 0.564 |
| Gender | (F_(1,51)_ = 1.680, *p* = 0.201 |
| ***Victim – annoyed*** | |
| Intention ⅹ Gender | (F_(1,51)_ = 0.034, *p* = 0.854 |
| Intention | (F_(1,51)_ = 41.559, *p* < 0.001, η_p_^2^ = 0.449 |
| Gender | (F_(1,51)_ = 1.294, *p* = 0.261 |
| ***Victim – feeling “bad”*** | |
| Intention ⅹ Gender | (F_(1,51)_ = 0.815, *p* = 0.371 |
| Intention | (F_(1,51)_ = 46.874, *p* < 0.001, η_p_^2^ = 0.479 |
| Gender | (F_(1,51)_ = 3.526, *p* = 0.066 |
| ***Moral Judgment Task*** | |
| ***Endorsement rate*** | |
| Taxonomy ⅹ Gender | (F_(2,100)_ = 1.523, *p* = 0.224 |
| Taxonomy | (F_(2,100)_ = 24.034, *p* < 0.001, η_p_^2^ = 0.325 |
| Gender | (F_(1,50)_ = 0.786, *p* = 0.380 |

**Table S5** Ages and questionnaire scores for subjects who completed the experimental protocol (n=162) in placebo (PLC), oxytocin (OXT) and vasopressin (AVP) groups (mean ± SD)

| **Measurement** | **PLC group**  **(n = 53)** | **OXT group**  **(n = 52)** | **AVP group**  **(n = 57)** | **F** | ***p*** |
| --- | --- | --- | --- | --- | --- |
| Age | 20.98±1.87 | 20.50±1.78 | 20.18±1.83 | 2.694 | 0.071 |
| Weschler verbal IQ | 112.33±4.87 | 114.15±5.60 | 114.02±5.53 | 1.918 | 0.15 |
| PANAS - positive (pre-test) | 27.92±6.72 | 29.71±6.84 | 27.79±5.82 | 1.466 | 0.234 |
| PANAS - negative (pre-test) | 17.87±5.83 | 19.96±6.67 | 18.61±6.65 | 1.444 | 0.239 |
| STAI - state (pre-test) | 41.30±11.10 | 42.58±10.88 | 41.3±9.72 | 0.258 | 0.773 |
| STAI - trait | 43.62±9.30 | 44.13±10.31 | 42.86±10.16 | 0.228 | 0.796 |
| ^a^PANAS - positive (post-test) | 27.42±7.97 | 27.60±7.43 | 26.68±6.73 | 0.238 | 0.789 |
| ^a^PANAS - negative (post-test) | 17.98±7.10 | 17.92±6.64 | 16.84±6.98 | 0.476 | 0.622 |
| ^a^STAI - state (post-test) | 39.96±11.21 | 41.12±11.21 | 40.19±11.04 | 0.157 | 0.855 |
| BDI | 10.06±9.16 | 10.63±9.23 | 10.11±8.68 | 0.067 | 0.935 |
| ASQ | 21.66±5.60 | 21.56±5.77 | 20.49±6.00 | 0.691 | 0.502 |
| NEO-FFI - agreeableness | 42.42±5.39 | 42.96±5.48 | 43.00±5.12 | 0.202 | 0.817 |
| NEO-FFI - conscientiousness | 40.04±6.63 | 41.02±5.94 | 38.28±5.87 | 2.793 | 0.064 |
| NEO-FFI - extraversion | 37.98±7.24 | 38.77±6.82 | 38.91±6.38 | 0.292 | 0.747 |
| NEO-FFI - neuroticism | 35.42±8.06 | 35.65±9.02 | 36.14±8.55 | 0.103 | 0.902 |
| NEO-FFI - openness | 42.96±4.80 | 42.87±5.70 | 42.84±5.60 | 0.008 | 0.992 |
| IRI - perspective taking | 11.87±4.37 | 11.92±4.69 | 12.33±3.89 | 0.192 | 0.825 |
| IRI - fantasy scale | 16.09±5.16 | 17.27±4.02 | 15.89±4.65 | 1.367 | 0.258 |
| IRI - empathy concern | 17.09±3.58 | 17.29±3.49 | 17.30±4.16 | 0.05 | 0.951 |
| IRI - personal distress | 9.58±4.19 | 9.15±4.95 | 9.33±4.45 | 0.12 | 0.887 |
| IRI total score | 54.64±11.56 | 55.63±10.38 | 54.86±10.73 | 0.121 | 0.886 |
| SQ - adaptive form | 11.85±2.98 | 12.29±2.77 | 12.46±2.69 | 0.672 | 0.512 |
| SQ - egocentric form | 5.70±2.68 | 6.17±2.77 | 6.54±2.74 | 1.321 | 0.27 |
| SQ - pathological form | 7.96±3.48 | 7.71±3.53 | 8.18±3.99 | 0.216 | 0.806 |
| SQ total score | 25.51±7.42 | 26.17±7.76 | 27.18±7.86 | 0.657 | 0.52 |
| EPQ - idealism | 58.42±12.76 | 59.37±13.59 | 57.75±13.32 | 0.203 | 0.817 |
| EPQ - relativism | 64.75±12.17 | 65.12±11.67 | 64.26±11.60 | 0.072 | 0.931 |
| CTQ - emotional abuse | 8.13±3.51 | 7.92±3.54 | 7.65±3.37 | 0.268 | 0.765 |
| CTQ - physical abuse | 6.08±1.72 | 6.17±2.40 | 5.98±2.06 | 0.115 | 0.892 |
| CTQ - sexual abuse | 5.75±1.81 | 5.87±1.61 | 5.40±1.41 | 1.232 | 0.294 |
| CTQ - emotional neglect | 10.19±4.51 | 9.96±3.66 | 10.05±4.09 | 0.041 | 0.96 |
| CTQ - physical neglect | 7.58±2.36 | 7.63±2.88 | 7.35±2.26 | 0.202 | 0.817 |
| CTQ total score | 37.74±10.63 | 37.56±10.32 | 36.44±9.92 | 0.259 | 0.772 |
| EMBU - mother - emotion warmth | 20.87±5.77 | 22.50±4.53 | 22.96±4.24 | 2.757 | 0.067 |
| EMBU - mother - over protection | 17.08±4.57 | 18.19±4.97 | 17.30±4.66 | 0.82 | 0.442 |
| EMBU - mother - rejection | 8.23±3.24 | 8.02±3.30 | 8.05±3.04 | 0.065 | 0.938 |
| EMBU - father - emotion warmth | 20.19±6.12 | 18.94±4.95 | 19.4±5.17 | 0.708 | 0.494 |
| EMBU - father - over protection | 15.51±4.52 | 15.48±3.86 | 15.18±3.80 | 0.115 | 0.892 |
| EMBU - father - rejection | 7.81±2.52 | 7.92±2.43 | 8.12±2.78 | 0.205 | 0.815 |
| ICS - horizontal collectivism | 49.51±8.51 | 50.17±8.30 | 47.23±8.52 | 1.845 | 0.161 |
| ICS - vertical collectivism | 48.47±8.87 | 50.08±6.98 | 46.14±9.52 | 2.925 | 0.057 |
| ICS - horizontal individualism | 53.30±7.78 | 54.60±7.16 | 53.35±7.95 | 0.487 | 0.616 |
| ICS - vertical individualism | 46.25±8.73 | 47.23±7.98 | 45.88±8.49 | 0.372 | 0.69 |

ASQ: Autism Spectrum Quotient; BDI: Beck’s Depression Inventory; CTQ: Childhood Trauma Questionnaire; EMBU: Parenting Style Scale (Egna Minnen av Barndoms Uppforstran); EPQ: Ethics Position Questionnaire; ICS: Individualism-Collectivism Scale; IQ: Intelligence Quotient; IRI: Interpersonal Relation Inventory; NEO-FFI: NEO Five-Factor Inventory; PANAS: Positive and Negative Affective Scale; SQ: Selfishness Questionnaire; STAI: State Trait Anxiety Inventory. Superscript Symbols ‘a’ denotes one female subject in PLC group excluded due to the failure of post-test data collection.

**Table S6.** Examples of scenarios in the Moral Judgment Task

**Non-moral**

*Standard Turnips*

You are a farm worker driving a turnip-harvesting machine. You are approaching two diverging paths.

By choosing the path on the left you will harvest ten bushels of turnips. By choosing the path on the right you will harvest twenty bushels of turnips. If you do nothing your turnip-harvesting machine will turn to the left.

Is it appropriate for you to turn your turnip-picking machine to the right in order to harvest twenty bushels of turnips instead of ten?

*Train or Bus*

You need to travel from New York to Boston in order to attend a meeting that starts at 2:00 PM. You can take either the train or the bus.

The train will get you there just in time for your meeting no matter what. The bus is scheduled to arrive an hour before your meeting, but the bus is occasionally several hours late because of traffic. It would be nice to have an extra hour before the meeting, but you cannot afford to be late.

Is it appropriate for you to take the train instead of the bus in order to ensure you not being late for your meeting?

*Coupons*

You have gone to a bookstore to buy $50 worth of books. You have with you two coupons.

One of these coupons gives you 30% off of your purchase price. This coupon expires tomorrow. The other coupon gives you 25% off your purchase price, and this coupon does not expire for another year.

Is it appropriate for you to use the 30%-off coupon for your present purchase so that you will have another coupon to use during the coming year?

**Moral Impersonal**

*Sculpture (3)*

You are visiting the sculpture garden of a wealthy art collector. The garden overlooks a valley containing a set of train tracks. A railway workman is working on the tracks, and an empty runaway trolley is heading down the tracks toward the workman.

The only way to save the workman's life is to push one of the art collector's prized sculptures down into the valley so that it will roll onto the tracks and block the trolley's passage. Doing this will destroy the sculpture.

Is it appropriate for you to destroy the sculpture in order to save this workman's life?

*Standard Trolley (1-3)*

You are at the wheel of a runaway trolley quickly approaching a fork in the tracks. On the tracks extending to the left is a group of five railway workmen. On the tracks extending to the right is a single railway workman. If you do nothing the trolley will proceed to the left, causing the deaths of the five workmen.

The only way to avoid the deaths of these workmen is to hit a switch on your dashboard that will cause the trolley to proceed to the right, causing the death of the single workman.

Is it appropriate for you to hit the switch in order to avoid the deaths of the five workmen?

*Vaccine Policy (4)*

You work for the Bureau of Health, a government agency. You are deciding whether or not your agency should encourage the use of a certain recently developed vaccine. The vast majority of people who take the vaccine develop an immunity to a certain deadly disease, but a very small number of people who take the vaccine will actually get the disease that the vaccine is designed to prevent.

All the available evidence, which is very strong, suggests that the chances of getting the disease due to lack of vaccination are much higher than the chances of getting the disease by taking the vaccine.

Is it appropriate for you to direct your agency to encourage the use of this vaccine in order to promote national health?

**Moral Personal**

*Footbridge (2)*

A runaway trolley is heading down the tracks toward five workmen who will be killed if the trolley proceeds on its present course. You are on a footbridge over the tracks, in between the approaching trolley and the five workmen. Next to you on this footbridge is a stranger who happens to be very large.

The only way to save the lives of the five workmen is to push this stranger off the bridge and onto the tracks below where his large body will stop the trolley. The stranger will die if you do this, but the five workmen will be saved.

Is it appropriate for you to push the stranger on to the tracks in order to save the five workmen?

*Submarine*

You are the captain of a military submarine travelling underneath a large iceberg. An onboard explosion has caused you to lose most of your oxygen supply and has injured one of your crew who is quickly losing blood. The injured crew member is going to die from his wounds no matter what happens. The remaining oxygen is not sufficient for the entire crew to make it to the surface.

The only way to save the other crew members is to shoot dead the injured crew member so that there will be just enough oxygen for the rest of the crew to survive.

Is it appropriate for you to kill the fatally injured crew member in order to save the lives of the remaining crew members?

*Sophie's Choice***

It is wartime and you and your two children, ages eight and five, are living in a territory that has been occupied by the enemy. At the enemy's headquarters is a doctor who performs painful experiments on humans that inevitably lead to death.

He intends to perform experiments on one of your children, but he will allow you to choose which of your children will be experimented upon. You have twenty-four hours to bring one of your children to his laboratory. If you refuse to bring one of your children to his laboratory he will find them both and experiment on both of them.

Is it appropriate for you to bring one of your children to the laboratory in order to avoid having them both die?

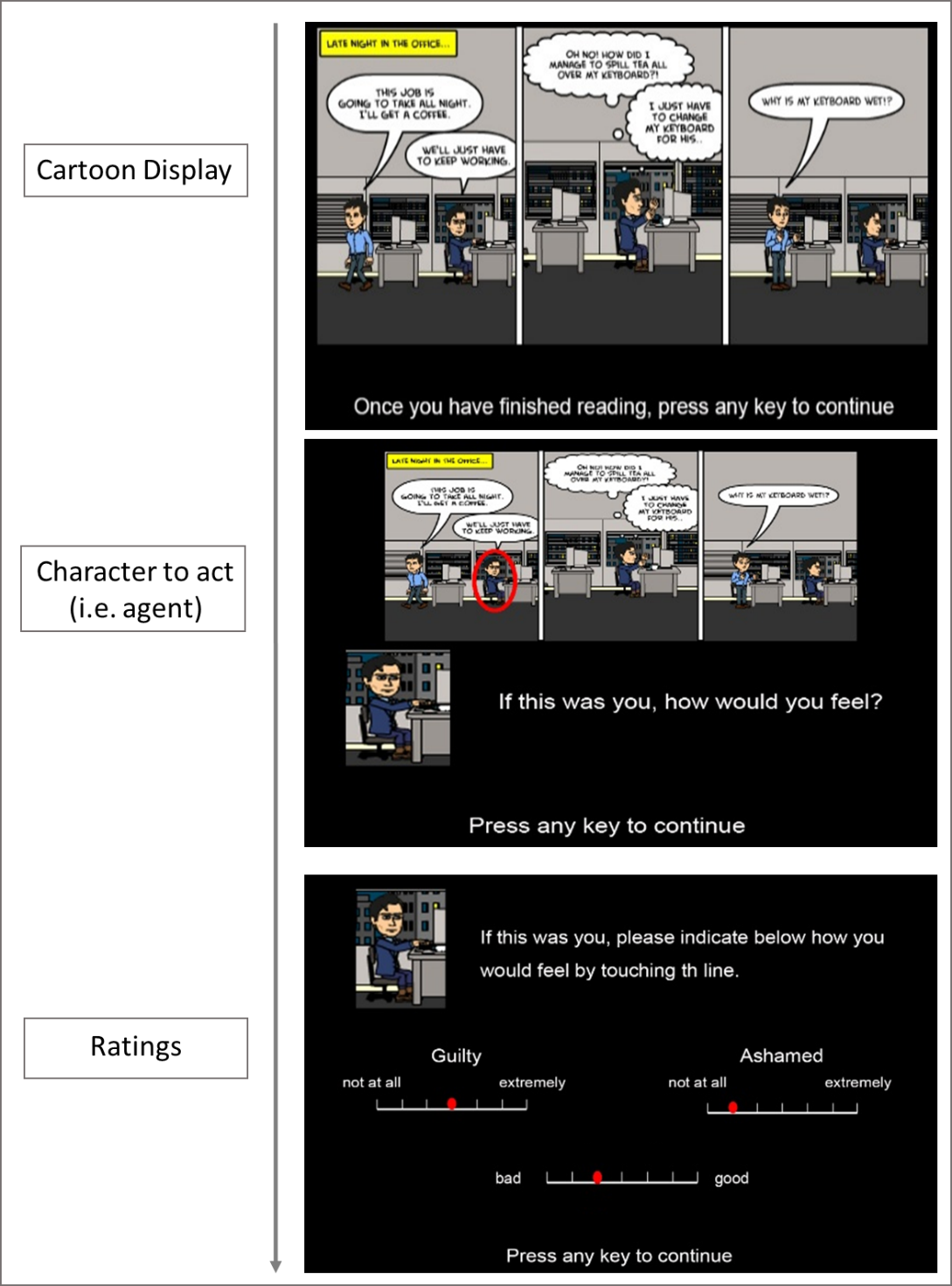
**Figure S1. Example of a single trial (e.g. Intention - deliberate harm; Character - agent) for the Moral Emotions Task.** Subjects were instructed to watch the presented cartoon silently and imagined the situations before they gave rating scores in terms of guilty, ashamed, feeling “bad” when acting as an agent and annoyed and feeling “bad” when acting as a victim. There was no time limitation for watching the cartoons or giving responses. For convenience the sentences in the figure are presented in English but for the actual experiment they were presented in Chinese.

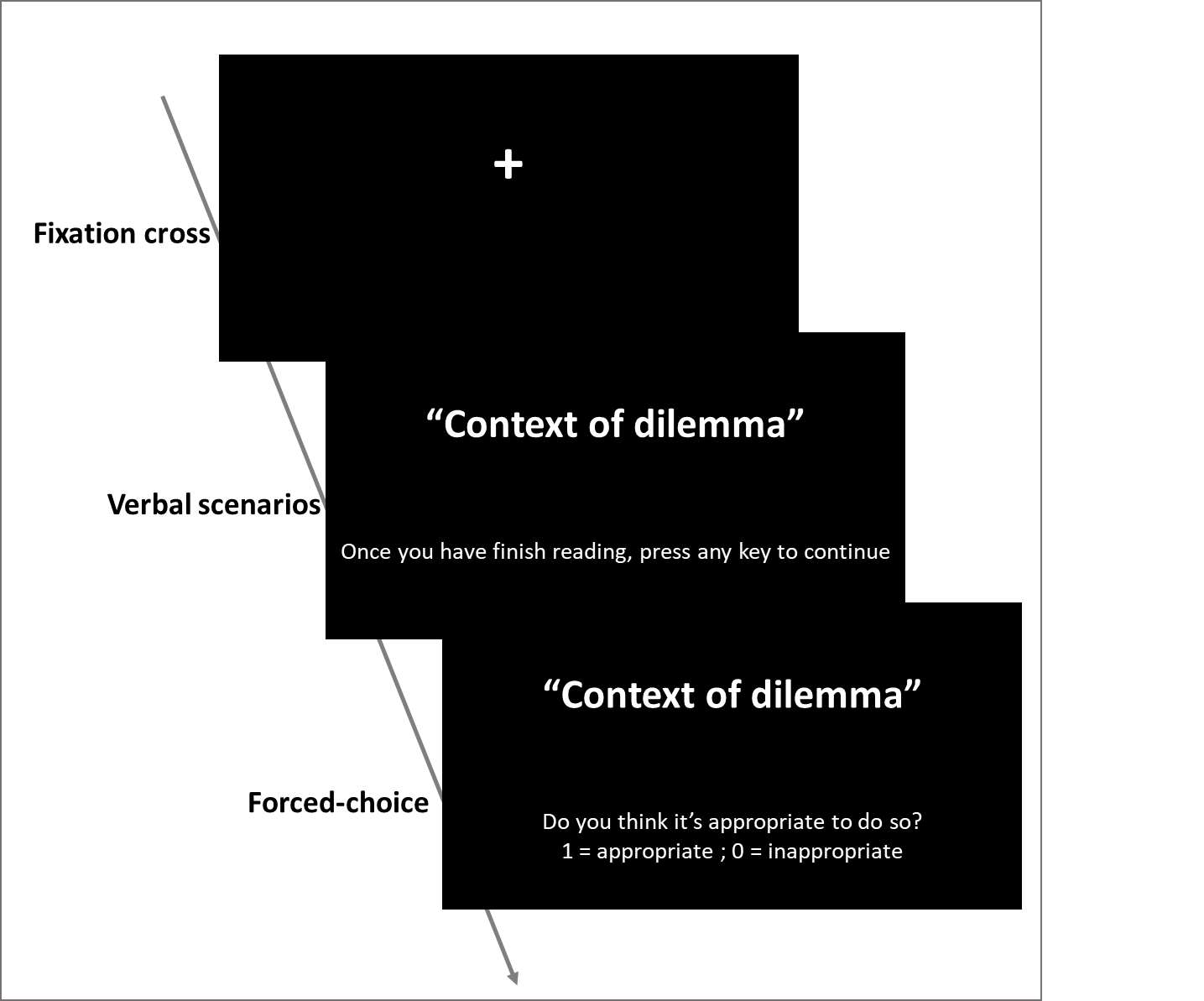

**Figure S2. Example of a single trial for the Moral Judgment Task.** Following a 1s fixation cross, each verbal description depicting a non-moral, moral impersonal, or moral personal scenario was randomly displayed. Subjects were instructed to read the descriptions silently and imagined the situations described in them before giving a forced choice towards the posed question to the current scenario. There was no time limitation for reading the description or giving responses. For convenience the sentences in the figure are presented in English but for the actual experiment they were presented in Chinese.

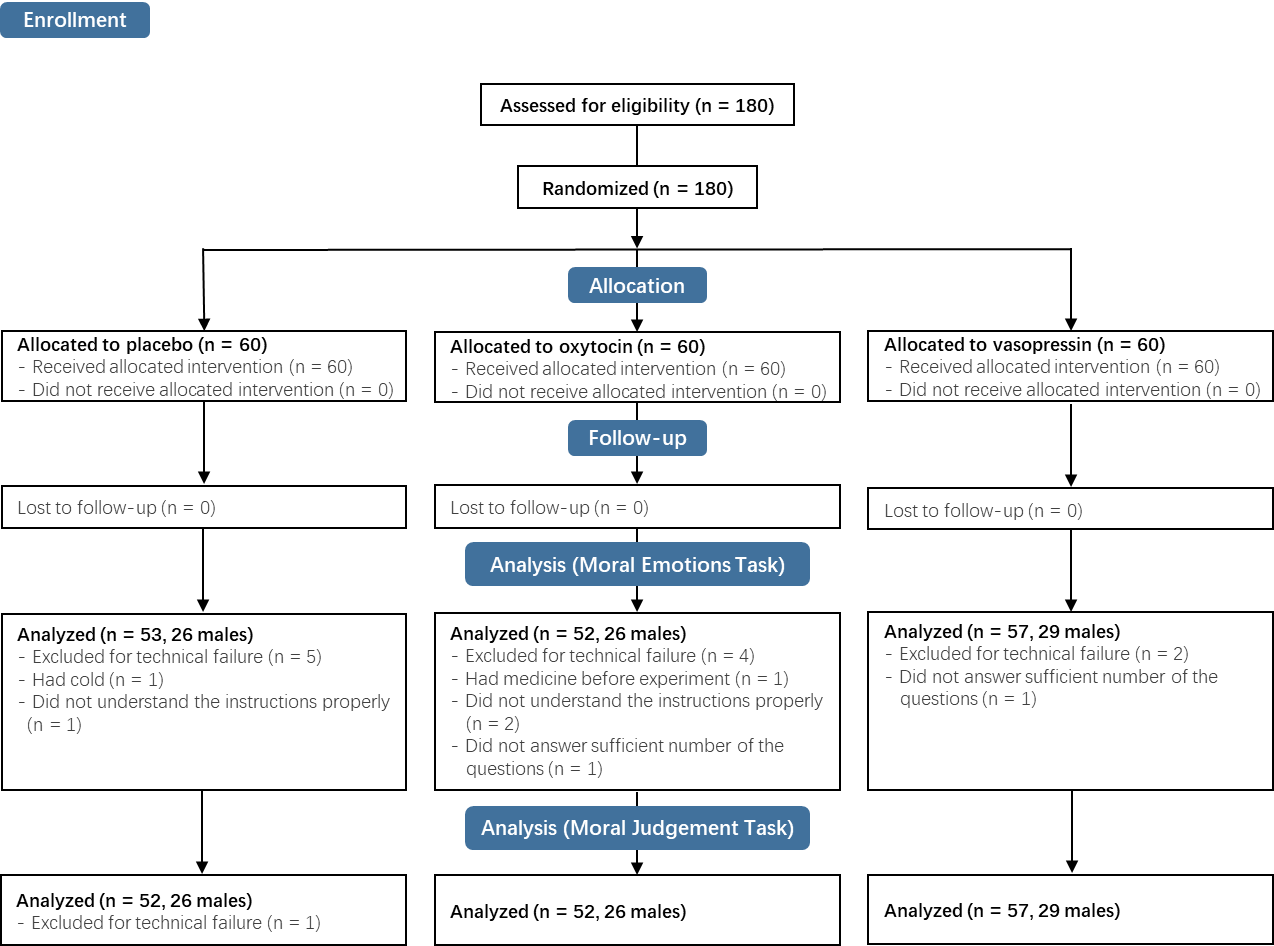
**Figure S3.** CONSORT flow chart displaying exclusion and rationale.
